# Upregulated toll-like receptors 2 and 4 expression underlies delayed diabetic wound healing

**DOI:** 10.1101/611053

**Authors:** Surabhi Bajpai, Ashish Dhayani, Praveen Kumar Vemula, Rakesh Mishra

## Abstract

Diabetes is marked by delayed wound healing response and prolonged phase of inflammation. Toll-like receptors (TLRs) are the key pathogen recognition receptors known to regulate inflammation in wound healing. Therefore, we aim to investigate the role of TLRs in delayed healing of diabetic wounds. TLR (1-9) expression studies were conducted using J774 macrophage cell line. Further, primary macrophages were isolated from wound tissues of *db/db* mice, subjected to RT-PCR and western blot analysis. The RT-PCR expression of TLRs revealed that expression of TLR2, TLR4, TLR5, TLR6, TLR7 and TLR8 were significantly upregulated in macrophage cells, cultured in high glucose medium. Expression of TLR 1-9 were significantly upregulated in wound tissues of *db/db* mice. However, TLR1, TLR2 and TLR4, TLR6, TLR7 were only found upregulated in the primary macrophages. Only TLR2 and TLR4 exhibited significantly increased protein expression. Further, TLR2 and TLR4 inhibition by OxPAPC (1-palmitoyl-2-arachidonyl-*sn*-glycero-3-phosphorylcholine) impregnated hydrogels, resulted in a significantly increased rate of wound healing in *db/db* mice via decreasing the significant levels of TNF-α and IL-1β. These results show that increased TLR 2 and TLR 4 expression underlies delayed wound healing in diabetes and a hydrogel impregnated with OxPAPC may be a promising therapeutic strategy for the management of diabetic wounds. Our study goes closer in understanding the molecular mechanism and provides a novel treatment method in diabetic wounds.

## Introduction

Diabetes is afflicting nearly 422 million people worldwide[1] and its prevalence is associated with increasing incidences of diabetes associated disorders. Type I and Type II diabetes both are marked by increased inflammation as evidenced by increased monocyte activity[2][3]. Inflammation is the key regulator in the diabetes-related disorders like delayed wound healing[4][5]. However, the cause of this prolonged inflammation in diabetes and underlying mechanisms are yet unknown. Therefore, a clearer understanding of the mechanisms underlying prolonged inflammation can be the key to success for designing targeting drugs for control of inflammation in diabetes.

Toll-like receptors (TLRs) are highly conserved pattern recognition receptors (PRRs) expressed in various cells involved in immune system like monocytes, neutrophils and macrophages[6], acting as key players in inflammation. A number of germline encoding pattern recognition receptors (PRRs) are known, of which, the TLR family is the major comprising of 13 members[7]. These PRRs recognize the pathogen-associated molecular patterns (PAMPs) often associated with the invading pathogens. Upon PAMPs recognition the PRRs trigger multitude intracellular signaling pathways in the host involving the active participation of adapter molecules, kinases and various transcription factors[8]. This process culminates with the activation of several cytokines, chemokines, cell adhesion molecules and immunoreceptors[9]. However, these TLRs can also recognize certain host factors as “threat signals” when they are present in the form of abnormal molecular complexes produced as a result of cellular stress[10].

TLRs have been found to play a major role in type1[11] and type2 diabetes[12]. Recent findings have shown that hyperglycemia induces a proinflammatory state with increased TLR 2, 4 expressions in human monocytes[13]. In another study conducted on type 1 diabetic patients with microvascular complications, TLR2 and TLR4 and its downstream signaling factors were found to be upregulated[14]. TLR3 and TLR5 were found to be upregulated in bone marrow-derived macrophages of non-obese diabetic mice[15]. TLR2 and TLR4 also mediated inflammation in diabetic nephropathy[16]. Understanding the importance of TLRs in inflammation, we next sought to understand the role of TLRs in diabetes. Till now most of the studies in diabetes focused on the role of TLR2 and TLR4 but this is for the first time that the role of other TLRs (1-9) have been analyzed in diabetic wound healing. Further, it is necessary to understand whether TLR inhibitor or inducer could help in healing the diabetic wounds. However, systemic circulation of these inducers/inhibitors would compromise the immune system. Therefore, we have used local application of hydrogel which releases TLR inhibitor/inducer in response to the inflammation.

Hydrogels are formed by weak, non-covalent interactions when small amphiphilic molecules self-assemble to form three dimensional matrices in an aqueous medium. Hydrogels are used for a variety of biomedical applications from localized drug[17] or biomolecule delivery[18] to encapsulation of mammalian cells[19], their maintenance and differentiation etc.[20]. These small molecules can be designed to have functional groups that respond to various physiological stimuli such as pH[21], various kinds of enzymes[22][23] ultrasound[24], and hypoxia[25]. We have used amphiphilic molecule triglycerolmonostearate (TGMS) here which can self-assemble in water to form hydrogels and encapsulate a variety of hydrophobic drugs. TGMS is a GRAS (generally recognized as safe) agent by US-FDA. An interesting aspect of TGMS is the presence of an ester group linking the hydrophilic and hydrophobic part that can be cleaved by over-expressed inflammatory enzymes such as various esterase and proteases. In a proof-of-concept study, we have shown encapsulation of an immunosuppressive agent, tacrolimus in TGMS hydrogels to prevent graft rejection in vascularized composite allotransplantation (VCA) in a rat model[22] and pig model[26]. It has been proved that the release of drug through TGMS is stimulus-dependent i.e. in response to the severity of inflammation. Due to the stimulus dependent response, the graft rejection could be prevented for >100 days compared to control animals. Hence, we chose TGMS-hydrogel system to deliver TLR2 and4inducerLPS-Ek (lipopolysaccharide from *E. coli* K-12) and inhibitor OxPAPC during the delayed wound repair/healing phases in diabetic animals. Our TGMS based system would ensure the release of these molecules only during the inflammatory stimulus along with a more localized delivery and minimal release in the absence of stimulus (i.e. enzymes). Therefore, making TGMS a very efficient system for delivery of therapeutics in chronic non-healing wounds. Moreover, OxPAPC if given through a systemic route might lead to systemic immune suppression and off-target effects. We have tried to avoid this problem by choosing a localized delivery system.

Therefore, this study shows that an abnormality in TLR expression leads to delayed healing of wounds in diabetes employing *db/db* mice model. Additionally, we have also shown that hydrogels impregnated with TLR 2 and TLR 4 inducer and inhibitor could potentially treat the non-healing diabetic wounds.

## Material and methods

### Cell Culture

J774 macrophage cells were obtained from National Centre for Cell Sciences, Pune, India. The *in vitro* studies were conducted on J774 macrophages (1×10^7^ cells/ml) cultured in 5 mmol/l (low glucose) and 30 mmol/l glucose (high glucose) for 24hrs. Cells cultured in 30 mmol/l mannitol were used as an osmotic control. Cell viability was >90%.

### Animals

All the experiments were performed on 5 to 6 weeks old female, inbred *db/db* mice on C57 BL/6J background with average weight of 26±1 gram. These were obtained from National Institute of Immunology (NII), New Delhi, India. The animals were maintained under controlled laboratory conditions. All animal care and experimental procedures described in this study were specifically approved by the Animal Ethics Committees of Indian Institute of Technology, Kanpur, India (CPCSEA Registration Number: 810/GO/ac/03/CPCSEA), protocol number: IITK/IAEC/2013/1011 and Banasthali Vidyapith, Rajasthan, INDIA (Registration Number: 574/GO/ReBi/S/02/CPCSEA;24.02.2002), protocol Number: BU/3432/16-17. All surgery was performed under odium pentobarbital anesthesia, and all efforts were made to minimize suffering. The mice had *ad libitum* access to food and water. The study was conducted with the prior approval of the institutional animal ethical committee. The mice homozygous for a spontaneous mutation of leptin receptor gene (*Lepr^db^*) became notably obese and showed elevated levels of glucose around 3-4 weeks. The wildtype, non-diabetic (ND) mice were used as controls.

### Wound creation

A single, full thickness, 6mm diameter excision wound was made at the superficial level on the mid-dorsum of each *db/db* mice and wild type mice. On third day, the wound tissues were collected and subjected to RT-PCR. All the *in vivo* and *in vitro* studies were performed in quadruplicate. Skin from *db/db* and wild type mice was kept as controls.

### Primary macrophage isolation from incision wound

The incision wounds were employed for procuring primary macrophages from wound area specifically. Incision wounds created by single incision (6mm) on the back of mice with circular polyvinyl alcohol (PVA) sterile sponges (6mm diameter), implanted subcutaneously in each pocket created by incision on the back of the mice; at the site of excision wounds. PVA sponges were harvested within 72hr post wound creation, the reason being that the inflammation stage lasts for 3 days. A single wound cell suspension was generated by repeated compression of sponges which was filtered through a 70μm nylon cell strainer to remove debris. For macrophage isolations, magnetic cell sorting was done using mouse anti-CD11b tagged micro-beads[27]. The macrophage cells were cultured for24hr in DMEM.

### RT-PCR

Total RNA was extracted from J774 macrophages cultured in low glucose, high glucose, mannitol after 24hr of culture. The total RNA was also extracted from primary macrophages (wound origin) after 72hr post isolation/culture, using TRIZOL method. 5μg of RNA from each sample was subjected to cDNA synthesis using standardized protocol[28] and further PCR amplification (RT-PCR) using TLR (1-9) specific primers as described earlier[29]. The *β*-actin gene was used as house-keeping control. Amplification was performed in a thermal cycler (Labnet, USA) programmed for 30 cycles of denaturation at 94°C for 1min., annealing at 55°C for 1min., and extension at 72°C for 45s. The whole program was preceded by initial denaturation at 95°C for 5min and ended with a final extension at 72°C for 10 minutes. For analysis of each reaction, the PCR products (10μl) were analyzed on 2 % agarose gel. Gels were photographed under UV illumination on a gel documentation system. The mRNA expression levels were analyzed by Image J Software and the relative expression was calculated by comparing the band intensity of various TLRs with the intensity of *β*-actin (house-keeping gene).

### Western Blotting

For western blotting, the proteins from primary macrophages of diabetic and wild type mice wounds were separated on SDS-PAGE and transferred on nitrocellulose membrane (MDI, India). After blocking with 5% non-fat dry milk powder, the membranes were incubated with TLR 2,4,6,7 (Cell Signaling Technology) specific primary antibody and secondary antibody sequentially. The immuno-reactive proteins positive for TLR antibody were visualized/detected by chemiluminescence documentation system (BioRad, U.S.A.). Finally signal intensities were measured using NIH Image J software.

### Preparation of TGMS hydrogels containing OxPAPC and LPS-Ek

TGMS was procured from A. K. Scientific, USA. Dimethlysulphoxide (DMSO) used for gel preparation was purchased from Sigma Aldrich (D2650). Autoclaved distilled water was used for all gel preparation procedures. OxPAPC and LPS-Ek were purchased from Invivo Gen, San Diego, USA. 10% (w/v) Triglycerolmonostearate (TGMS) was weighed in a glass vial. To this, 20% (v/v) DMSO was added. The mixture was heated up to 60-70°C to dissolve and allowed to stand. Upon cooling, the mixture was re-heated to dissolve, and remaining amount of water was added to form uniform solution which was taken into 1ml syringe and allowed to stand for 3-4 days before use in animals. In the formulation of gels loaded with OxPAPC and LPS-Ek; 400μg/ml and 50μg/ml of respective amounts were dissolved in 200μl of DMSO and formulated into 1ml gels as described above. Further, OxPAPC (TLR 2,4 inhibitor) and lipopolysaccharide from *E. coli* K-12(LPS-Ek) (TLR 2, 4 inducer) was encapsulated in triglycerolmonostearate self-assembled hydrogels and applied on excision wounds of *db/db* mice.

### Wound Contraction

Hydrogels (100μl) loaded with OxPAPC and LPS-Ek were applied on the excision wounds of four groups of mice: Non diabetic treated with OxPAPC (ND OxPAPC), diabetic treated with OxPAPC (*db/db* OxPAPC), Non diabetic treated with lipopolysachharide (ND LPS), diabetic treated with lipopolysachharide (*db/db* LPS) with 4 animals in each group respectively. Wounds of *db/db* placebo, ND placebo groups were filled with hydrogels only whereas one *db/db* and non-diabetic group of controls was left untreated. Topical application continued twice a day at regular intervals till day 14. Gel was spread uniformly to cover the wound bed of mice and further applied at regular intervals. After receiving one treatment, each mouse was individually housed and fed *ad libitum*. Digital camera was used to record the wound pictures of each mouse. Wound diameter was measured every day manually by using a metric scale. Rate of wound closure (%) = [(Original wound area – Open wound area on final day)/ Original wound area] X 100[30].

### Cytokine assays

After the application of hydrogels (100μl) loaded with OxPAPC and LPS-Ek along with controls the tissue supernatants were collected on day 14. The secretion levels of TNF-α, IL-1β were detected using a commercially available enzyme-linked immunosorbent assay (ELISA) system (Raybiotech, US) following kit instructions.

### Statistical analysis

The data were analyzed by a two tailed unpaired student’s test using Graph Pad software. The *p*-value less than 0.05 were considered significant. All the experiments were performed in triplicate and data were represented as mean ± standard deviation.

## Results

### Gene expression of TLR increases in cultured J774 macrophage cell line

The RT-PCR expression pattern of TLR1-9 revealed that expression of TLR2, TLR4, TLR5, TLR6, TLR7, TLR8 were significantly upregulated (*p*<0.001) in macrophage cells cultured in high glucose medium as compared to the macrophages cultured in low glucose medium and mannitol (Fig1). Comparing the relative expression pattern from cells cultured in the presence of mannitol with cells cultured in high glucose, it can be concluded that the increase in expression of most of the TLRs was due to high glucose and not because of an osmotic effect.

**Fig 1.**
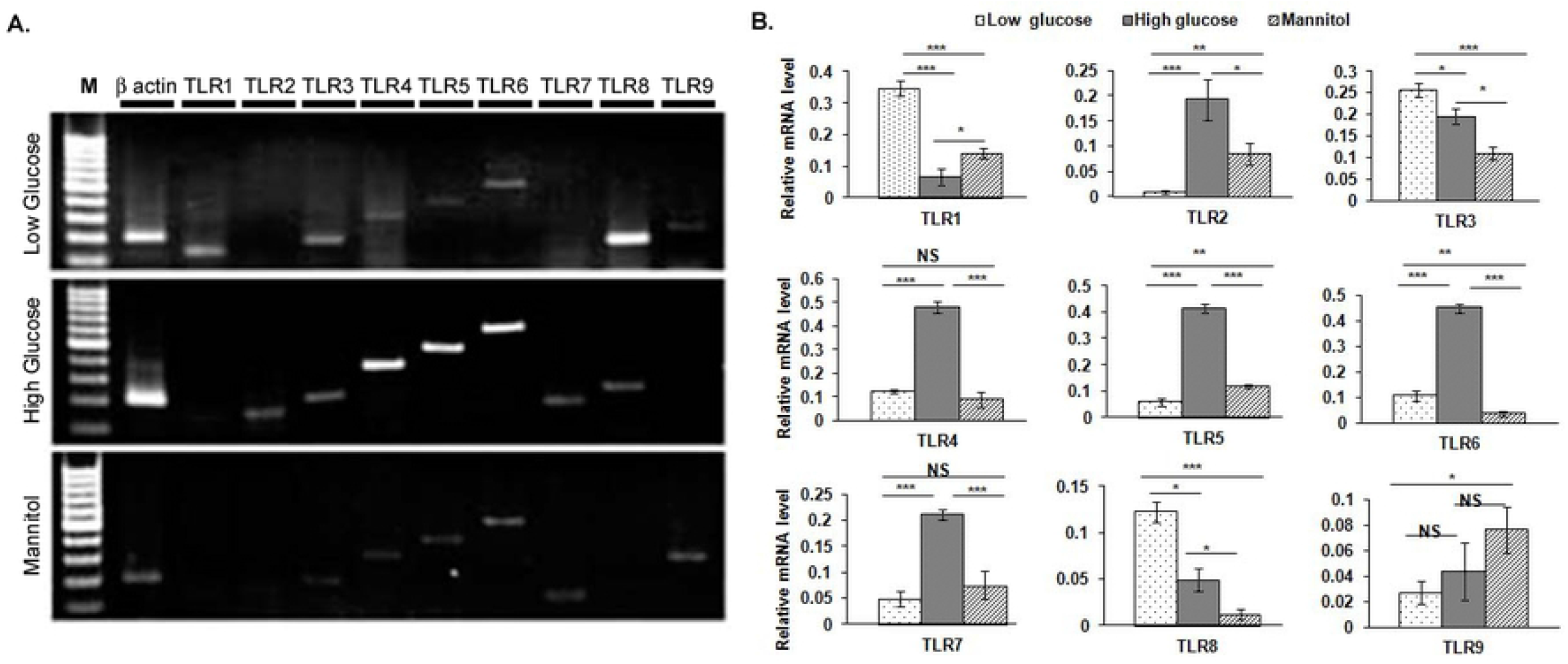
Increased expression of TLRs (2,4,5,6,7,8) in high glucose background. Expression of various TLRs from J774 macrophages (Fig 1A) cultured in low glucose (5mmol/l) and high glucose (30mmol/l) and taking mannitol (30mmol/l) as osmotic control. Semiquantitative analysis (Fig 1B) of expression levels of different TLRs in J774 macrophages shows that TLR2, TLR4, TLR5, TLR6, TLR7, TLR8 are significantly upregulated (*p*<0.001) in macrophage cells cultured in high glucose medium as compared to the macrophages cultured in low glucose medium and mannitol, as osmotic control. *β*-actin used as internal control. Marker-100 bp ladder, statistical significance: *p*<0.05*, *p*<0.01**, *p*<0.001***.

### Gene expression of TLR increases in diabetic skin and wound concurrently

The results of the RT-PCR from skin and wound tissues of *db/db* mice exhibited significantly increased (*p*<0.001) TLRs (1,2,3,4,5,6,7,9) expression as compared to the un-injured skin and wound tissues of wild type mice as control. TLR8 also exhibited increased expression in diabetic skin (*p*<0.05) and wound (*p*<0.01) tissues as compared to the skin and wound tissues of wild-type mice. Importantly, there was no significant change (p>0.05) in the level of different TLR(1-9) in the *db/db* wound tissue as compared to the un-injured *db/db* skin which implicates that these TLRs were increased at a systemic level in diabetic mice and not restricted to the wound area specifically (Fig 2). This suggested us that probably the systemic upregulation in the level of TLRs in diabetes leads to prolonged inflammation resulting in aberrant healing at the localized site of the wound where TLR-expressing macrophages accumulate.

**Fig 2.**
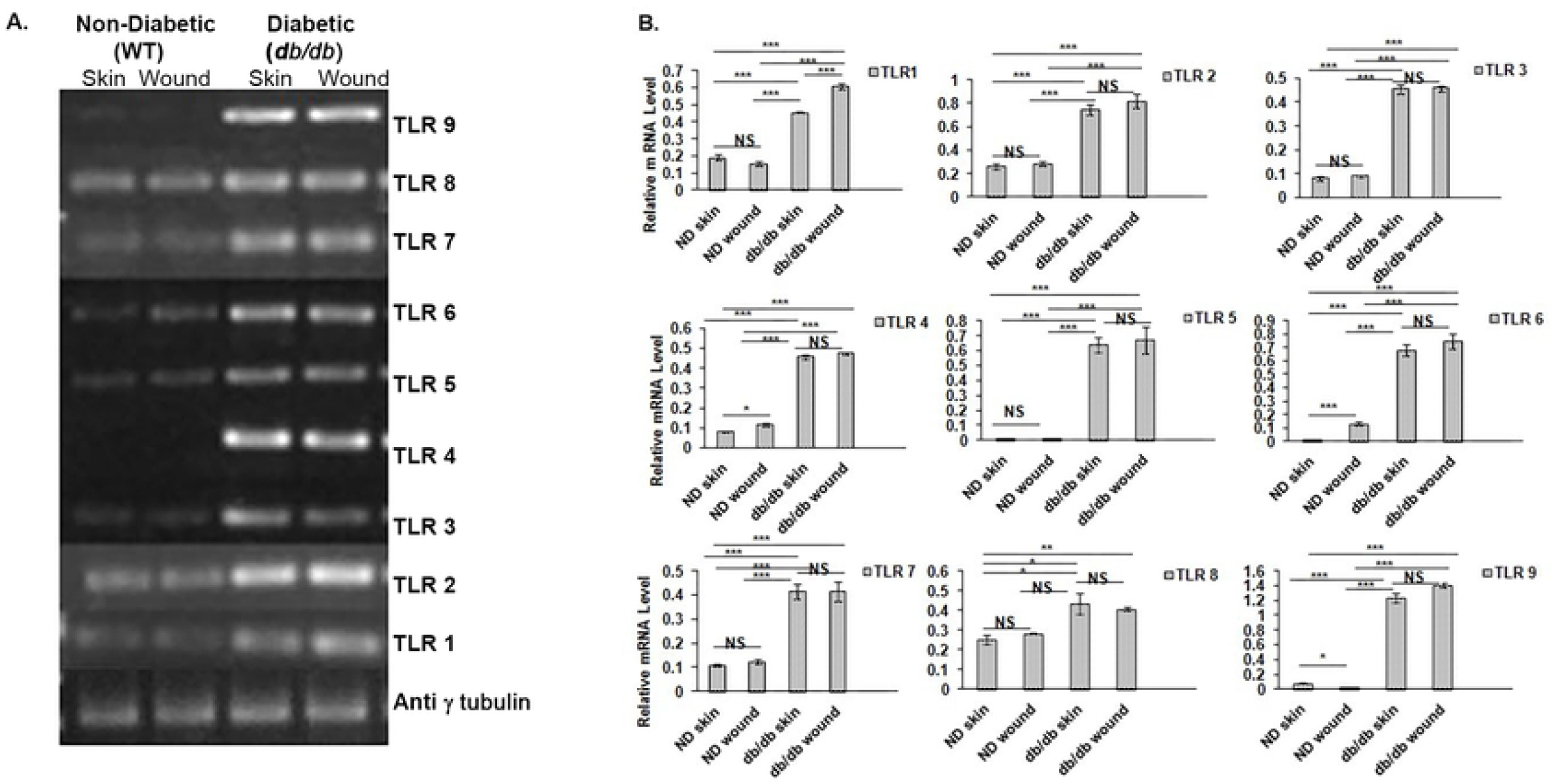
Enhanced expression of different TLRs in un-injured skin and wound tissues of wild type and *db/db* mice. Semiquantitative analysis of expression level of different TLRs as observed in agarose gel (A) are analyzed (B). There is a significant increase in levels of TLRs (1-9) in skin and wound tissues of *db/db* mice as compared to the control (un-injured skin and wound tissues of wild type mice). *β*-actin was used as internal control. Statistical significance: *p*<0.05*, *p*<0.01**, *p*<0.001***.

### Gene expression of TLR increases in diabetic wounds

Expression of TLR 2-9 were found significantly upregulated (*p*<0.001) in wound tissues of *db/db* mice as compared to wild type control (Fig 3).

**Fig 3.**
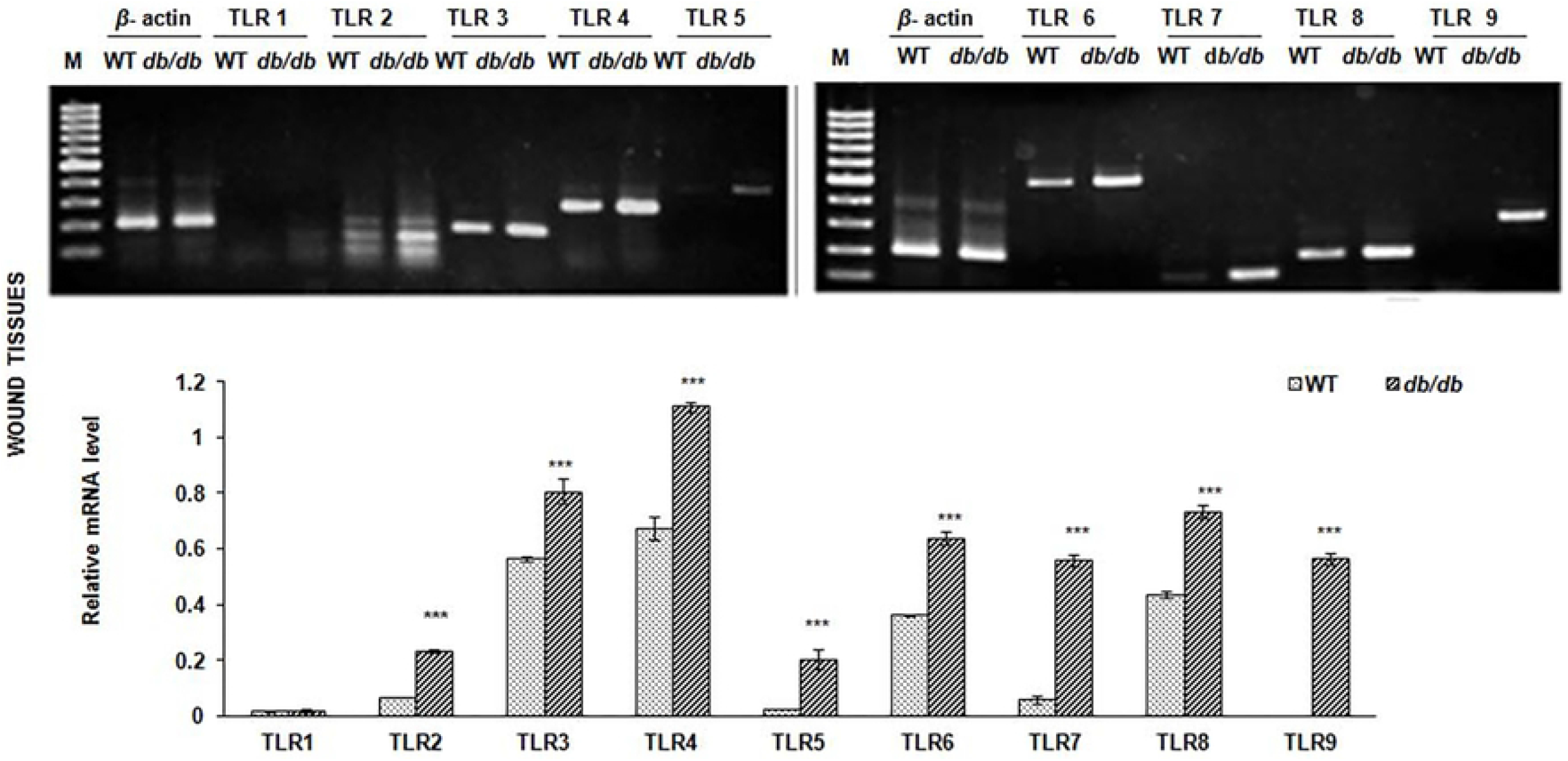
Upregulation of TLRs (2-9) in wounds of *db/db* mice. Semiquantitative expression of various TLRs as observed in wound tissues of wild type and *db/db* mice suggests upregulation of TLR 2-9 in wound tissues of *db/db* mice. *β*-actin used as internal control. 100bp DNA ladder used as marker. Statistical significance: *p*<0.05*, *p*<0.01**, *p*<0.001***.

### Gene expression of TLR increases in cultured primary macrophages

TLR1, TLR2 (p<0.05) and TLR4, TLR6, TLR7 (*p*<0.001) were only found upregulated in the primary macrophages derived from wound tissues of *db/db* mice as compared to wild type (Fig 4).

**Fig 4.**
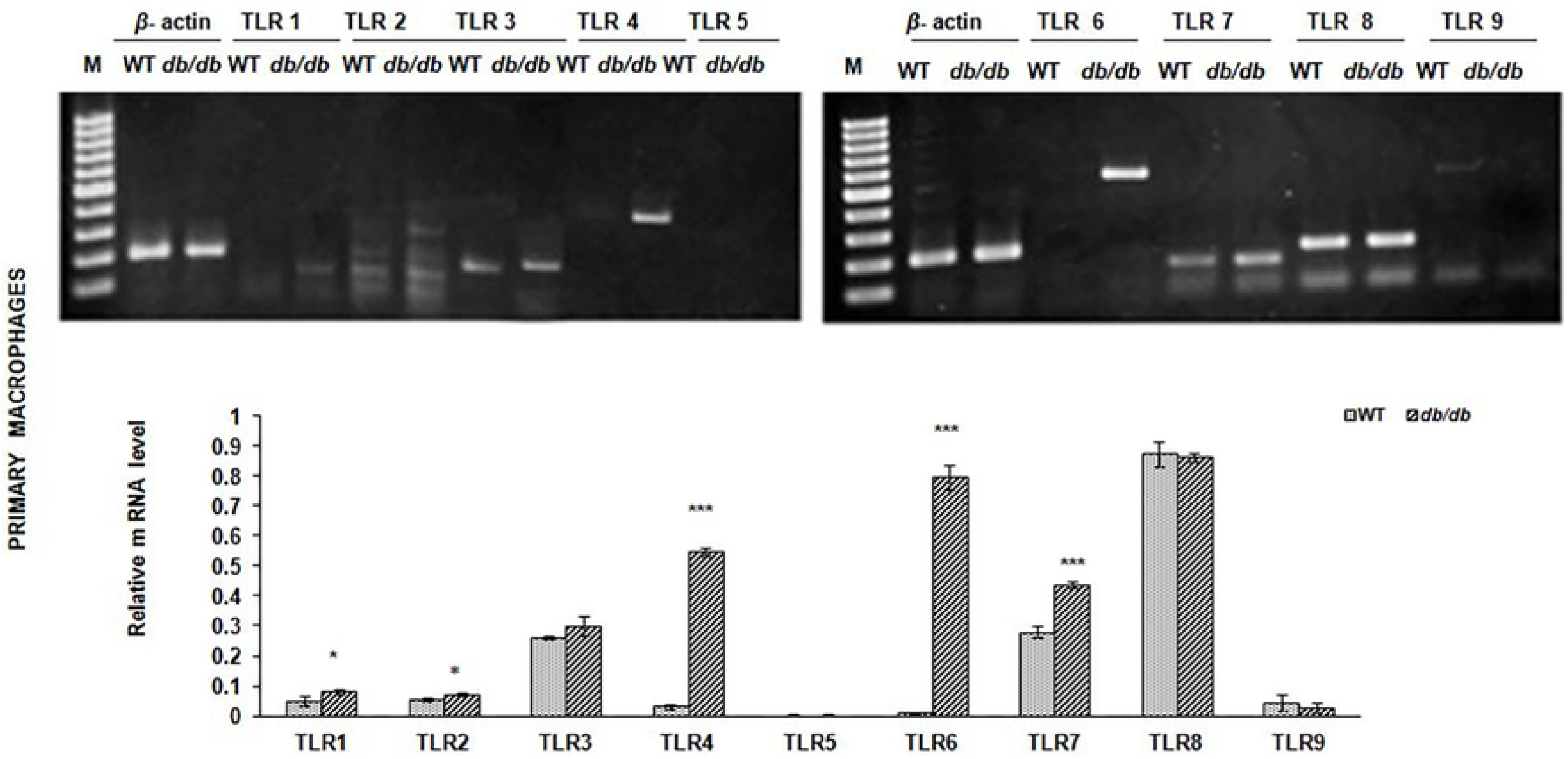
Increased RNA expression of various TLRs in *db/db* mice primary macrophages (wound origin). Semiquantitative analysis done using primary microphages cultured from wound tissues collected from *db/db* mice showed significant increase in TLR1 & TLR2 (p<0.05) and also in TLR4, TLR6 and TLR7 ((*p*<0.001). *β*-actin was used as internal control. Marker-100bp DNA ladder. Statistical significance: *p*<0.05*, *p*<0.01**, *p*<0.001***.

### TLR protein upregulation in cultured primary macrophages

Out of the all TLRs expressed at the transcription level in primary macrophages from *db/db* mice only TLR2,4,6,7 were found to be significantly upregulated (*p*<0.001) at the protein level as well (Fig 5). This implicates that the TLR proteins are being expressed at the protein level at the wound site where macrophages are attracted as a result of the injury. Probably, downstream signaling from this increased level of TLR resulted into the enhanced production of inflammatory cytokines. This would further implicate that there could be an increased number of macrophages which could result in the increased TLR 2,4,6,7 protein expression.

**Fig 5.**
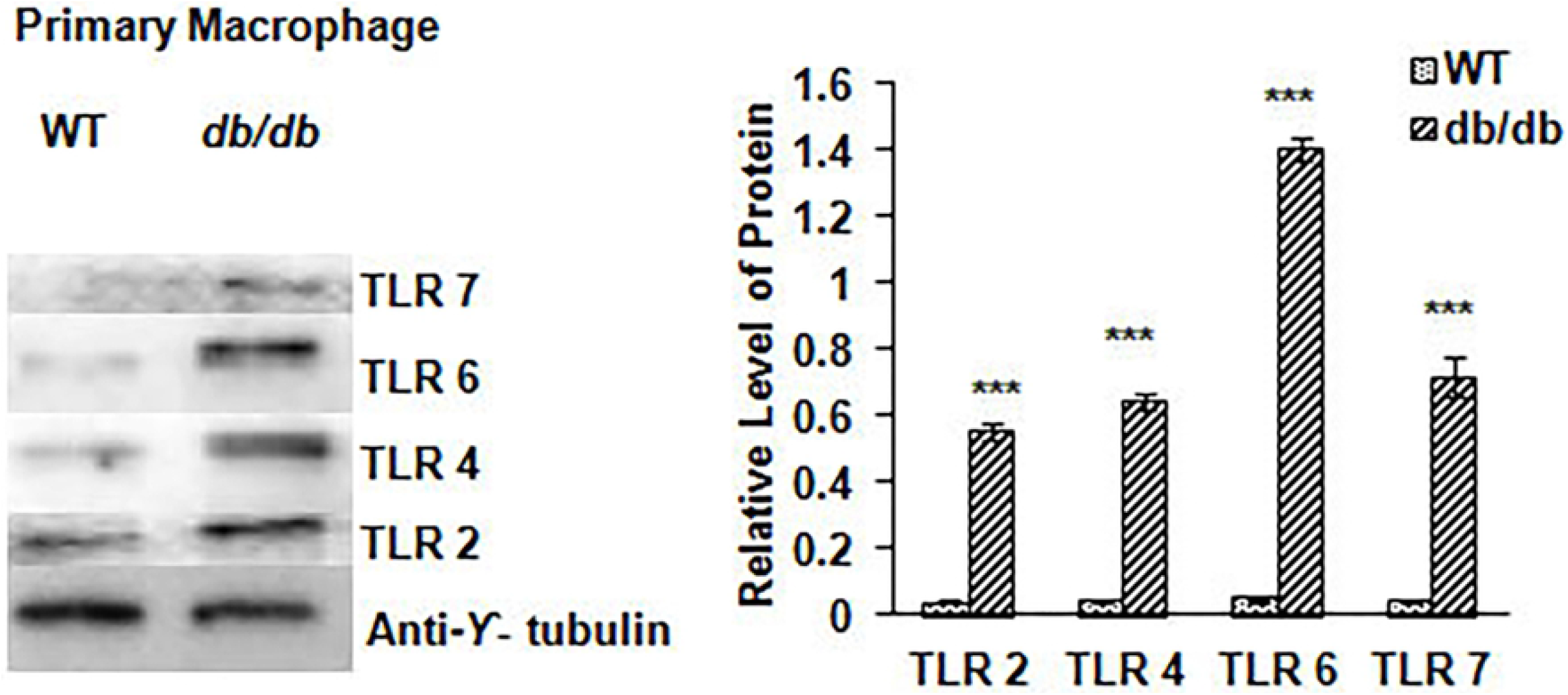
Upregulated TLR protein expression in primary macrophages isolated from *db/db* mice. Except TLR1, having upregulated RNA expression, other TLRs (2,4,6 and 7), having upregulated RNA, show corresponding significant upregulation (*p*<0.001) at the protein level as well Anti-*γ*-tubulin used as internal control. Semi quantitative analysis of protein level of different TLRs using NIH Image J software. Statistical significance: *p*<0.05*, *p*<0.01**, *p*<0.001***.

### Hydrogels impregnated with OxPAPC promoted wound contraction in *db/db* mice

The results showed significantly greater wound contraction in the hydrogel impregnated with OxPAPC *db/db* group at 7 days after treatment compared with the control, placebo groups (hydrogel) and hydrogel+LPS groups (*p* < 0.001, n = 4). After fourteen days post wound creation, the rate of wound closure in diabetic mice was 88.88 ± 2.92% in the hydrogel + OxPAPC group, whereas 44.43 ± 4.62%, 72.5 ± 6.67%, and 61.1 ± 4.21% in the *db/db* control, placebo and hydrogel+LPS groups of mice, respectively. However, significantly fast wound contraction (*p* 0.001, n = 4) was reported in the control non diabetic group of mice compared with the placebo groups, hydrogel+OxPAPC group and hydrogel+LPS groups at day 14 post wound creation (Fig 6).

**Fig 6.**
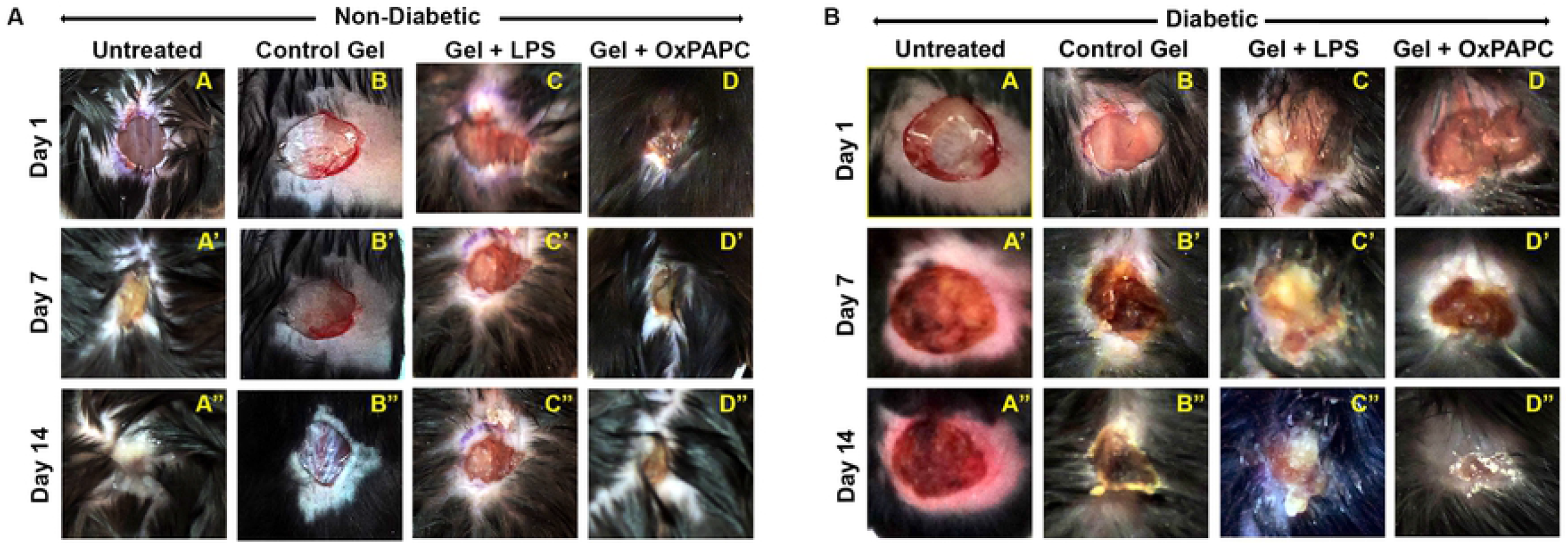
Faster wound contraction with hydrogel impregnated OxPAPC in diabetic mice. This figure illustrates wound contraction in various groups of mice observed on day 1, day 7, day 14 post treatment with different hydrogels. Two groups formed: non-diabetic (A) and diabetic (B). Differrent mice from non-diabetic are further grouped as non diabetic mice (untreated), non diabetic mice control (hydrogel treated /placebo), non diabetic mice treated with hydrogel+LPS, non diabetic mice treated with hydrogel+OxPAPC. And similarly from diabetic group, the subgroups are *db/db* mice, *db/db* mice placebo (hydrogel treated/placebo), *db/db* mice treated with hydrogel+LPS, *db/db* mice treated with hydrogel+ OXPAPC. *db/db* mice treated with hydrogel impregnated with OxPAPC exhibited greater degree of wound closure. Wound diameter was recorded by Nikon digital camera.

### Hydrogels loaded with OxPAPC decreased the cytokine level at wound site

Significant decrease in the level of TNFα and IL-1β (*p*<0.001) was observed in *db/db* mice treated with hydrogel + OxPAPC as compared to the controls, placebo and hydrogel+LPS treated groups of mice. However, the result was found to be less significant in all four groups of non-diabetic mice (*p*>0.05) (Fig 7).

**Fig 7.**
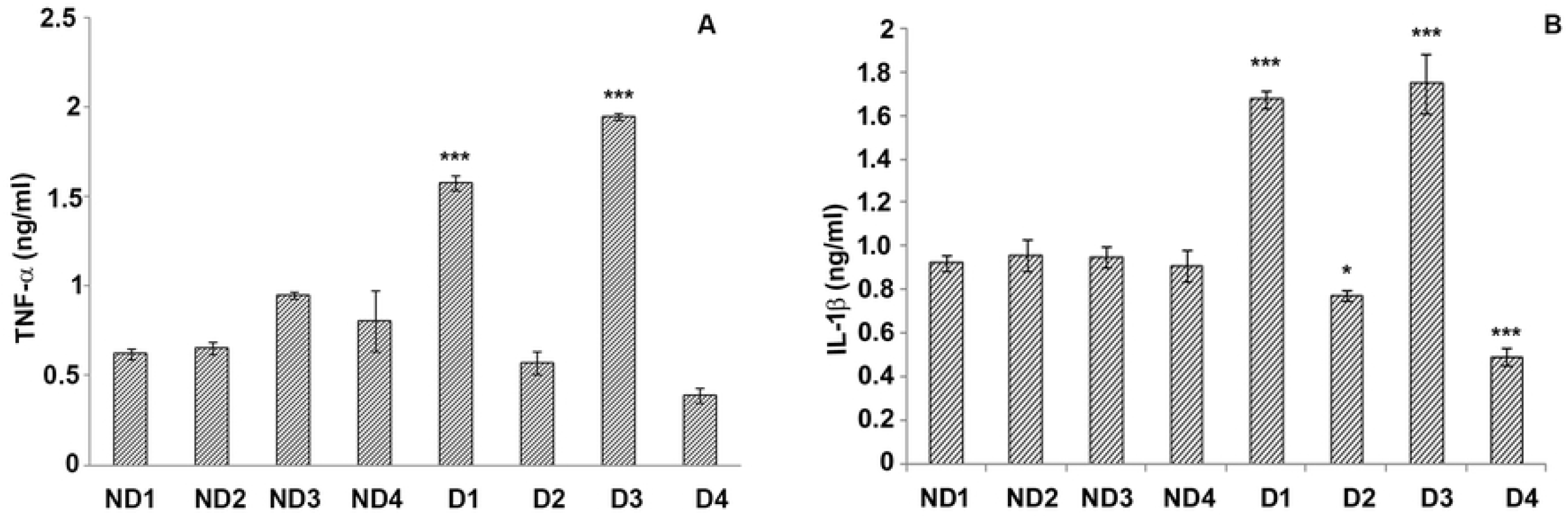
Diabetic mice treated with hydrogel + OxPAPC showed decreased level of TNFα and IL-1β. The levels of cytokines, TNF-α (A) and IL-1β (B), in different groups of mice estimated by ELISA. The groups includes: ND1(non diabetic mice, control), ND2 (non diabetic mice treated with placebo /hydrogel only), ND3 (non diabetic mice treated with LPS+hydrogel), ND4 (non diabetic mice treated with OxPAPC+ hydrogel), D1(Diabetic *db/db* mice), D2(Diabetic *db/db* mice treated with placebo/hydrogel only), D3 (Diabetic *db/db* mice treated with LPS+hydrogel), D4 (Diabetic *db/db* mice treated with OXPAPC+ hydrogel). Statistical significance: *p*<0.05*, *p*<0.01**, *p*<0.001***.

## Discussion

Persistent inflammation is the hallmark of diabetes[3][30]. This inflammation in diabetes is associated with an increase in the level of several inflammatory cytokines like interleukin-1β, TNF-α and chemokines[31]. TLRs and its downstream signaling is the main source of production of these proinflammatory cytokines. TLR 2 and 4 are majorly known to be involved in the pathogenesis of diabetes and its associated disorders[13][14][16]. Our studies were based on the hypothesis that overexpression of TLR genes might play a significant role in delayed wound healing process in diabetic patients. In our studies, we have shown these conclusively using *in vitro* and *in vivo* based systems.

The results of this study show that TLR2, TLR4, TLR5, TLR6, TLR7, TLR8 were significantly upregulated (*p*< 0.001) in J774 macrophage cells cultured in high glucose medium which mimics the diabetic condition *in vitro*. This implicates that hyperglycemia in diabetes may lead to an upregulation in expression of TLRs in the macrophages. These results are in accordance with the previous work conducted using human monocytes (THP-1) cell line, where increased glucose leads to increased TLR 2, 4 expression[13]. These results were further confirmed in our lab by gene expression studies conducted on primary macrophages isolated from wound tissues of *db/db* mice which again confirmed the upregulation of TLR1, TLR2and TLR4, TLR6, TLR7 (*p*<0.001). These results for the very first time show the involvement of many TLRs (TLR 2-8) altogether in hyperglycemic condition.

The next we wanted to study whether the increased level of TLRs could also alter the wound healing in diabetes. It is well evident that hyper inflammation in diabetic wounds causes excessive production of proinflammatory cytokines leading to homing of inflammatory cells which elicits pro-apoptotic genes and damages re-epithelization, often resulting in wound matrix degradation [32]. This study also showed that diabetic wound exhibited an increased level of TLRs (TLR 1-9) as compared to the skin and wound of wildtype mice altogether. Importantly, out of all TLRs expressed at the transcript level in primary macrophages from *db/db* mice only TLR 2 and 4 were found to be significantly upregulated (*p*<0.001) at the protein level as well. This result is in accordance with the results of a study conducted on Type-2 diabetic patients where TLR 2, 4were found to be increased in monocytes[12]. The mechanism behind the systemic increase in TLR 2 and 4 expressions can be via PKC-α and PKC-δ by activating NADPH oxidase[13]. This increase in TLR expression could be attributed towards increased inflammation thereby causing a delayed healing in diabetic mice. TLR either through MyD88 dependent or through MAL/TRIF/TRAM pathway (or both in case of TLR4) lead to the activation of MAPKK and IKK complex resulting into activation and nuclear translocation of AP-1 and NF-kβ from macrophages which promote the secretion of multiple inflammatory cytokines downstream [33].

To understand the wound healing response better and to know whether TLR2,4 downregulation may heal the diabetic wounds in optimum time, we decided to encapsulate and deliver OxPAPC, an inhibitor of TLR2,4 to see if it helps in better/faster wound healing by modulating the pro-inflammatory response and alleviating inflammation. We envisioned using an inflammation-responsive hydrogel-based system to deliver the modulator (OxPAPC) locally on wounds of *db/db* mice to prevent its effects on the systemic immune system. In our study for the first time, we have shown an accelerated rate of wound closure upon the topical use of OxPAPC loaded hydrogels in a wound model using *db/db* mice, *in vivo*. Further, our results clearly indicate the significant decrease in the level of pro-inflammatory cytokines IL-1β and TNF-α upon topical application of hydrogels encapsulating OxPAPC over wounds of diabetic mice. The encapsulation of TLR inhibitor into TGMS gels led to sustained and inflammation responsive release of the cargo at the wound site, thus aiding in better wound closure compared to placebo. Therefore, these OxPAPC impregnated hydrogels can be implicated as a novel targeting therapy for targeting site-specific early closure of diabetic wounds without compromising the immune system systemically, but this needs further investigations.

Summarizing this data, we can state that systemic upregulation in TLR2 and TLR4 expression causes delayed wound healing in diabetic mice. The use of hydrogels loaded with TLR2 and TLR4 inhibitor could ameliorate the delayed wound closure in diabetic mice targeting the wound site without compromising the immune system.

## Acknowledgements

The author SB acknowledges Prof S. Ganesh, Lab 5, Department of Biological Sciences and Bioengineering, Indian Institute of Technology, Kanpur and Department of Bioscience and Biotechnology, Banasthali Vidyapith, Rajasthan as hosts for DST INSPIRE project. SB is also thankful to Dr. Nagrajan at National Institute of Immunology (NII), New Delhi for providing genetic diabetic mice (*db/db*) model. AD would like to thank University Grants Commission for Senior Research Fellowship. PV thanks Department of Biotechnology (DBT) for the Ramalingaswamy re-entry fellowship. SB is thankful to Dr. Nagrajan at National Institute of Immunology, New Delhi. RM is thankful to Indian Council for Medical Research (ICMR) for Senior Research Fellowship.

## References

1. Global report on diabetes. World Health Organisation. 2016;1–84.

2. Devaraj S, Glaser N, Griffen S, Wang-Polagruto J, Miguelino E, Jialal I. Increased monocytic activity and biomarkers of inflammation in patients with type 1 diabetes. Diabetes. 2006;55: 774–779. doi:10.2337/diabetes.55.03.06.db05-1417

3. Donath MY. Targeting inflammation in the treatment of type 2 diabetes: Time to start. Nat Rev Drug Discov. Nature Publishing Group; 2014;13: 465–476. doi:10.1038/nrd4275

4. Wetzler C, Kampfer H, Stallmeyer B, Pfeilschifter J, Frank S. Large and sustained induction of chemokines during impaired wound healing in the genetically diabetic mouse: Prolonged persistence of neutrophils and macrophages during the late phase of repair. J Invest Dermatol. 2000;115: 245–253. doi:10.1046/j.1523-1747.2000.00029.x

5. Ebaid H, Ahmed OM, Mahmoud AM, Ahmed RR. Limiting prolonged inflammation during proliferation and remodeling phases of wound healing in streptozotocin-induced diabetic rats supplemented with camel undenatured whey protein. BMC Immunol. BMC Immunology; 2013;14: 1. doi:10.1186/1471-2172-14-31

6. O’Mahony DS, Pham U, Iyer R, Hawn TR, Liles WC. Differential constitutive and cytokine-modulated expression of human Toll-like receptors in primary neutrophils, monocytes, and macrophages. Int J Med Sci. 2008;5: 1–8. doi:10.7150/ijms.5.1

7. Takeda K, Akira S. Toll-like receptors. Curr Protoc Immunol. 2015;109: 1–10. doi:10.1007/978-3-319-29785-9_2

8. Akira S, Uematsu S, Takeuchi O. Pathogen recognition and innate immunity. Cell. 2006;124: 783–801. doi:10.1016/j.cell.2006.02.015

9. Akira S, Takeda K. Toll-like receptor signalling. Nat Rev Immunol. 2004;4: 499–511. doi:10.1038/nri1391

10. Beg AA. Endogenous ligands of Toll-like receptors: Implications for regulating inflammatory and immune responses. Trends Immunol. 2002;23: 509–512. doi:10.1016/S1471-4906(02)02317-7

11. D. Z. Toll like receptors and Type I Diabetes [Internet]. Advances in experimental medicine and biology. 2010. doi:10.1007/978-90-481-3271-3

12. Dasu MR, Devaraj S, Park S, Jialal I. Increased Toll-Like Receptor (TLR) activation and TLR ligands in recently diagnosed type 2 diabetic subjects. Diabetes Care. 2010;33: 861–868. doi:10.2337/dc09-1799

13. Dasu MR, Devaraj S, Zhao L, Hwang DH, Jialal I. High glucose induces toll-like receptor expression in human monocytes Mechanism of activation. Diabetes. 2008;57: 3090–3098. doi:10.2337/db08-0564

14. Devaraj S, Jialal I, Yun J-M, Bremer A. Demonstration of Increased TLR2 and TLR4 Expression in Monocytes of Type 1 Diabetic Patients with Microvascular Complications. Metabolism. 2011;60: 256–259. doi:10.1016/j.metabol.2010.01.005. Demonstration

15. Mohammad MK, Morran M, Slotterbeck B, Leaman DW, Sun Y, von Grafenstein H, et al. Dysregulated Toll-like receptor expression and signaling in bone marrow-derived macrophages at the onset of diabetes in the non-obese diabetic mouse. Int Immunol. 2006;18: 1101–1113. doi:10.1093/intimm/dxl045

16. Mudaliar H, Pollock C, Komala MG, Chadban S, Wu H, Panchapakesan U. The role of Toll-like receptor proteins (TLR) 2 and 4 in mediating inflammation in proximal tubules. Am J Physiol Physiol. 2013;305: F143–F154. doi:10.1152/ajprenal.00398.2012

17. Vemula PK, Cruikshank GA, Karp JM, John G. Self-assembled prodrugs: An enzymatically triggered drug-delivery platform. Biomaterials. Elsevier Ltd; 2009;30: 383–393. doi:10.1016/j.biomaterials.2008.09.045

18. Loebel C, Rodell CB, Chen MH, Burdick JA. Shear-thinning and self-healing hydrogels as injectable therapeutics and for 3D-printing. Nat Protoc. 2017;12: 1521–1541. doi:10.1038/nprot.2017.053

19. Chen S, Shi J, Zhang M, Chen Y, Wang X, Zhang L, et al. Mesenchymal stem cell-laden anti-inflammatory hydrogel enhances diabetic wound healing. Sci Rep. 2015; doi:10.1038/srep18104

20. Hwang NS, Varghese S, Theprungsirikul P, Canver A, Elisseeff J. Enhanced chondrogenic differentiation of murine embryonic stem cells in hydrogels with glucosamine. Biomaterials. 2006; doi:10.1016/j.biomaterials.2006.06.033

21. Ling Y, Lu M. Preparation and Characterization of pH and Temperature Dual Responsive-, Poly(N-isopropylacrylamide-co-itaconic acid) Hydrogels Using DMF and Water as Mixed Solvents. Polym J. 2008;40: 592–600. doi:10.1295/polymj.PJ2007213

22. Gajanayake T, Olariu R, Leclère FM, Dhayani A, Yang Z, Bongoni AK, et al. A single localized dose of enzyme-responsive hydrogel improves long-term survival of a vascularized composite allograft. Sci Transl Med. 2014;6. doi:10.1126/scitranslmed.3008778

23. Purcell BP, Lobb D, Charati MB, Dorsey SM, Wade RJ, Zellars KN, et al. Injectable and bioresponsive hydrogels for on-demand matrix metalloproteinase inhibition. Nat Mater. 2014;13: 653–661. doi:10.1038/nmat3922

24. Huebsch N, Kearney CJ, Zhao X, Kim J, Cezar CA, Suo Z, et al. Ultrasound-triggered disruption and self-healing of reversibly cross-linked hydrogels for drug delivery and enhanced chemotherapy. Proc Natl Acad Sci. 2014;111: 9762–9767. doi:10.1073/pnas.1405469111

25. Yu Jicheng, Zhang Yuqi, Ye Yanqi, DiSanto Rocco, Sun Wujin, Ranson Davis, Ligler Frances S., Buse John B. ZG. Microneedle-array patches loaded with hypoxia-sensitive vesicles provide fast glucose-responsive insulin delivery. Proc Natl Acad Sci. 2015;112: 8260–8265.

26. Fries CA, Lawson SD, Wang LC, Slaughter K V., Vemula PK, Dhayani A, et al. Graft-implanted, enzyme responsive, tacrolimus-eluting hydrogel enables long-term survival of orthotopic porcine limb vascularized composite allografts: A proof of concept study. PLoS One. 2019;14: e0210914. doi:10.1371/journal.pone.0210914

27. Khanna S, Biswas S, Shang Y, Collard E, Azad A, Kauh C, et al. Macrophage dysfunction impairs resolution of inflammation in the wounds of diabetic mice. PLoS One. 2010;5. doi:10.1371/journal.pone.0009539

28. Singh PK, Singh S, Ganesh S. The Laforin-Malin Complex Negatively Regulates Glycogen Synthesis by Modulating Cellular Glucose Uptake via Glucose Transporters. Mol Cell Biol. 2011;32: 652–663. doi:10.1128/mcb.06353-11

29. Sodhi A, Tarang S, Kesherwani V. Concanavalin A induced expression of Toll-like receptors in murine peritoneal macrophages in vitro. Int Immunopharmacol. 2007;7: 454–463. doi:10.1016/j.intimp.2006.11.014

30. Donath MY, Shoelson SE. Type 2 diabetes as an inflammatory disease. Nat Rev Immunol. Nature Publishing Group; 2011;11: 98–107. doi:10.1038/nri2925

31. Shanmugam N, Reddy MA, Guha M, Natarajan R. High glucose-induced expression of proinflammatory cytokine and chemokine genes in monocytic cells. Diabetes. 2003;52: 1256–1264. doi:10.2337/diabetes.52.5.1256

32. Loots MAM, Lamme EN, Zeegelaar J, Mekkes JR, Bos JD, Middelkoop E. Differences in cellular infiltrate and extracellular matrix of chronic diabetic and venous ulcers versus acute wounds. J Invest Dermatol. Elsevier Masson SAS; 1998;111: 850–857. doi:10.1046/j.1523-1747.1998.00381.x

33. Takeda K, Akira S. Toll-like receptors in innate immunity. Int Immunol. 2005;17(1):1–14. doi:10.1093/intimm/dxh186

